# Leveraging the mass balances of cellular metabolism to infer absolute concentrations from relative abundance metabolomics data

**DOI:** 10.1101/2021.05.13.444095

**Authors:** Justin Y. Lee, Mark P. Styczynski

## Abstract

**Motivation:** As the large-scale study of metabolites and a direct readout of a system’s metabolic state, metabolomics has significant appeal as a source of information for many metabolic modelling platforms and other metabolic analysis tools. However, metabolomics data are typically reported in terms of relative abundances, which precluding use with tools where absolute concentrations are necessary. While chemical standards can be used to determine the absolute concentrations of metabolites, they are often time-consuming to run, expensive, or unavailable for many metabolites. A computational framework that can infer absolute concentrations without the use of chemical standards would be highly beneficial to the metabolomics community.

**Results:** We have developed and characterized MetaboPAC, a computational strategy that leverages the mass balances of a system to infer absolute concentrations in metabolomics datasets. MetaboPAC uses a kinetic equations approach and an optimization approach to predict the most likely response factors that describe the relationship between absolute concentrations and their relative abundances. We determined that MetaboPAC performed significantly better than the other approaches assessed on noiseless data when at least 60% of kinetic equations are known *a priori*. Under the most realistic conditions (low sampling frequency, high noise data), MetaboPAC significantly outperformed other methods in the majority of cases when 100% of the kinetic equations were known. For metabolomics datasets extracted from systems that are well-studied and have partially known kinetic structures, MetaboPAC can provide valuable insight about their absolute concentration profiles.

## Introduction

Since its inception, metabolomics has been used in a wide variety of applications, including identification of disease biomarkers, disease diagnosis, and even drug development [1, 2]. While genomics, proteomics, and transcriptomics provide an upstream view of cellular function and information about what may occur in a system, metabolomics is a direct readout of a system’s current metabolic state and has emerged as an important area of study for understanding what is actually occurring in a system [3]. As the systems-scale study of metabolites, metabolomics has the potential to be integrated into metabolic modelling frameworks to better understand how cellular systems function and react to endogenous or exogenous perturbations [4]. In order to develop accurate metabolic models, an ample amount of metabolomics data will be necessary.

Depending on the scope of a study and the metabolites of interest, researchers commonly use three analytical techniques to measure metabolomics: nuclear magnetic resonance (NMR) spectroscopy and gas and liquid chromatography-mass spectrometry (GC-MS and LC-MS, respectively). One of the key advantages of using NMR is that it is non-destructive [5], meaning samples can be reused. However, compared to GC-MS and LC-MS, NMR is significantly less sensitive, which is why many researchers turn to mass spectrometry when measuring hundreds or thousands of metabolites at low concentrations in a single sample [6]. LC-MS is able to detect more metabolites than GC-MS and does not require any sample derivatization [5, 7], but GC-MS still remains popular among researchers due to its relative inexpensiveness compared to other methods [5, 8].

The greatest weakness of these mass spectrometry approaches is quantification [9], as the metabolomics data resulting from using these instruments are typically relative abundances and not absolute concentrations. These relative abundances still allow for some types of analysis, including principal component analysis (PCA) [10] and t-tests, as relative abundance measurements of the same analyte can be compared from sample to sample. However, comparing the relative abundances of different metabolites has no meaning [11]. Even if two metabolites have similar absolute concentrations, their peaks on a chromatogram and therefore their relative abundances can be radically different because of how chemicals with different structures and properties are derivatized, ionized, or fragmented [12, 13]. Similarly, peaks with comparable intensities do not necessarily imply equal absolute concentrations. This precludes the use of raw metabolomics data in many computational tools used to study metabolism. For example, several metabolic modelling platforms, such as MetDFBA, TMFA, and LK-DFBA [14–16] can directly integrate metabolite data into their frameworks, but they all require absolute concentrations.

Many researchers use chemical standards in mass spectrometry to quantify metabolites. However, these standards can be costly, time consuming to use, and unavailable for certain metabolites [12, 17, 18]. Using standards may only be feasible for quantifying a few metabolites, but for the purposes of untargeted metabolomics, where one attempts to measure all metabolites [19], it quickly becomes infeasible. Untargeted metabolomics data is restricted to being used with only the most exploratory computational tools, like PCA, for semi-quantitative analysis [11]. A method for determining absolute concentrations without the use of chemical standards would expand the usability of metabolomics data in computational tools and would be incredibly beneficial to the metabolomics community.

As one of the most critical challenges preventing metabolomics from being more readily used in wider applications, there have previously been efforts attempting to quantify metabolomics data without chemical standards. Much of this work has focused on predicting ionization efficiencies of different chemicals, which is directly linked to the relative abundance output of a chromatogram in mass spectrometry. It has been shown that intrinsic thermodynamic properties, electrokinetic properties, structural properties, and solvent factors are all key factors that contribute to the prediction of ionization efficiencies [20, 21]. Recently, Liigand et al. developed a method for predicting ionization efficiencies using random forest machine learning [22]. Another recent approach is MetabQ, a calibration curve-free method for quantification of polar metabolites [23]. While MetabQ still requires chemical standards, they only need to be used once in the lifetime of an instrument to determine the relationship between relative abundances and absolute concentrations.

In this work, we have developed a new computational framework for inferring the most likely absolute concentrations from relative abundance metabolomics data for cellular metabolism, which we have named Metabolomics Prediction of Absolute Concentrations (MetaboPAC). MetaboPAC attempts to avoid the need for chemical standards by leveraging the mass balances of a metabolic system and determining the most biologically likely metabolic profiles. To the best of our knowledge, this is the first computational platform for standard-free inference of absolute concentrations using metabolic mass balances. MetaboPAC could play a significant role in improving the ability to readily integrate metabolomics data with metabolic modelling and other metabolic analysis tools in the future.

## Methods

### Synthetic models

To assess MetaboPAC on different types of possible metabolic systems, we created two synthetic models. The first synthetic model (Figure S1A) contains four metabolites and five fluxes, where the initial influx is a known, constant reaction rate. With four metabolites and four unknown fluxes, the system is determined, which allows for the fluxes to be trivially solved using Equation 1, where 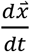 is the vector of the change in concentration over time for each metabolite, *S* is the stoichiometric matrix of the system, and 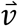 is the vector of fluxes. Since a determined system typically has a single unique flux profile solution, this simplification makes it a prime candidate for initial testing.

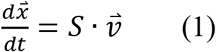

The second synthetic model (Figure S1B) contains four metabolites and eight fluxes. Once again, the influx is assumed to have a known, constant reaction rate. Unlike the determined model, the second model contains more unknown fluxes than metabolites and is therefore underdetermined. Furthermore, we have included two allosteric regulatory interactions, inhibiting flux v_3_ and promoting flux v_8_. Most biological systems are underdetermined and include metabolite-dependent, making this system a more complex and more relevant test for MetaboPAC. Both the determined and underdetermined with regulation synthetic models were constructed using Michaelis-Menten kinetics to model each reaction.

### Biological models

While synthetic models are pragmatic for initially developing and testing MetaboPAC, they do not sufficiently resemble biological models to allow generalization of initial results to later applications. To further evaluate the robustness of MetaboPAC, we examined models of *Escherichia coli* [24] and *Saccharomyces cerevisiae* [25] metabolism. Both of these systems are underdetermined and include numerous allosteric regulatory interactions, with the *E. coli* model containing 18 metabolites and 48 fluxes, and the *S. cerevisiae* model containing 22 metabolites and 24 fluxes. The kinetic reaction equations for both models include a mixture of Michaelis-Menten, Hill, and mass action kinetics.

### Response Factors

To emulate relative abundance data, we generated 20 sets of response factors for each metabolite found in the four systems evaluated in this work. Response factors describe the relationship between relative abundances and their absolute concentrations. Each response factor was randomly selected from a uniform distribution between 1 and 1000. These sets of response factors (*RF_T_*) were multiplied by the true absolute concentration values simulated by the kinetic models to calculate the relative abundances, assuming there is a direct linear relationship between the two (Equation 2). To infer absolute concentrations, the relative abundances are divided by the response factors predicted by MetaboPAC. The absolute concentrations for the systems used in this work ranged from 1e-4 mM to 20 mM.

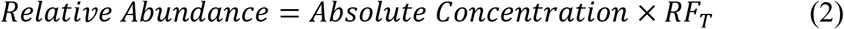

### Kinetic equations approach

If the kinetic rate law of each reaction in the system has been previously determined, the mass balances of the system and dynamic nature of time course metabolomics data can be leveraged to identify the response factors necessary to infer absolute concentrations. Based on the mass balances, the rate of change in concentration of a metabolite must always equal the sum of stoichiometrically balanced influxes and effluxes of the metabolite (Equation 1). When the kinetics of the reaction fluxes are known, each influx and efflux can be represented by a mathematical term containing kinetic parameters and the concentration of the metabolite(s) that participate in the reaction, either as a substrate or an allosteric regulator. Because only relative abundances and not absolute concentrations are available, the metabolite concentrations in these kinetic equations are replaced by their respective relative abundances divided by a response factor. The rate of change can be determined by calculating the difference in relative abundance at two subsequent timepoints and dividing by the change in time. Once again, the relative abundances in these rate of change calculations are divided by a response factor to infer absolute concentrations.

Across different timepoints in the metabolomics dataset, the response factors should remain constant for each metabolite, as they are not expected to change throughout an experiment. Together, the mass balances at each timepoint create a system of non-linear equations. A non-linear least-squares solver can determine the set of response factors that minimizes mass balance violations. These systems of equations must be determined or overdetermined to use the non-linear least-squares solver; as the number of timepoints in the data increases, the chance of having an underdetermined system of equations decreases. In the kinetic equations approach, this system of non-linear equations is solved 48 times (chosen based on the maximum number of local workers (12 workers, each used 4 times) when performing parallel computations) with different initial seeds selected from a uniform distribution and the medians of the predicted response factors are calculated at the conclusion of all the runs as the most likely set of response factors.

### Optimization approach

It is not uncommon for the kinetic equations of a reaction to be unknown, especially if the reaction is not in the most studied pathways of metabolism, such as central carbon metabolism. Instead of relying completely on the mass balances of the system to determine response factors, the optimization approach creates a minimization problem to predict the most likely set of response factors. In addition to minimizing mass balance violations in Equation 1 (without the known kinetic equations of the fluxes), there are also several penalties that can be added to the objective function to help identify sets of response factors that are biologically likely (Table S1). These penalties eliminate sets of response factors that lead to absolute concentrations that are biologically infeasible. For example, if a metabolite is the sole substrate of an enzyme, we expect the reaction rate to increase as the concentration of the metabolite increases. If this interaction is not observed between the inferred absolute concentration of the metabolite and the flux, the set of response factors would be heavily penalized. As in the kinetic equations approach, the optimization approach is performed 48 times with different initial seeds and the medians of the predicted response factors from all the runs is calculated to determine the most likely set of response factors.

### Combining the kinetic equations and optimization approaches

In many cases, the kinetic equations of the reactions in a system are only partially known. In this scenario, the kinetic equations and optimization approaches can be used in serial. First, the kinetic equations approach is used for the metabolite mass balance equations where all the kinetic equations of the influxes and effluxes are known. If only a few of the kinetic equations of the fluxes in a mass balance are known, it cannot be used in the kinetic equations approach; this can be a common occurrence when only a small percentage of the kinetic structure of the system is known. Only the response factors associated with the metabolites present in the useable mass balances can be identified in this step. After predicting all the possible response factors using the kinetic equations approach, the optimization approach proceeds as described above, except the response factors that have already been identified are fixed within the optimization problem and the remaining response factors are predicted. A workflow for this process is presented in Figure 1.

**Figure 1.**
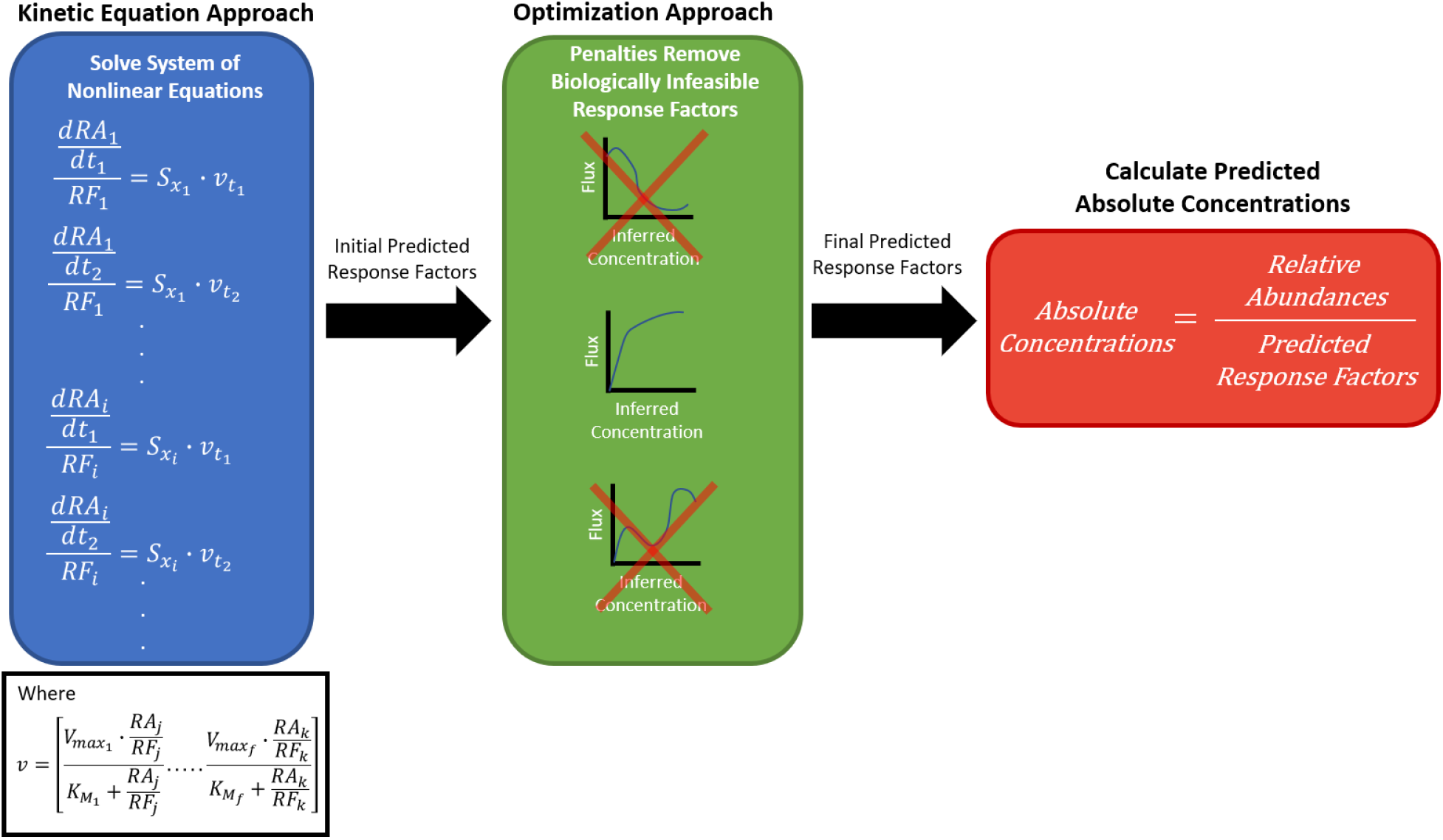
MetaboPAC workflow for inferring absolute concentrations from relative abundances in metabolomics datasets. In the kinetic equations approach, the mass balances at each timepoint are used to create a system of non-linear equations where the response factors in the useable mass balances are predicted. These initial predicted response factors are transferred and fixed in the optimization approach, where penalties are used to eliminate possible sets of the remaining response factors. The final predicted response factors are used to infer the absolute concentrations of the data. *RA_i_* is the relative abundance and *RF_i_* is the unknown response factor of the *i*th metabolite, *t_n_* is a particular timepoint in the data, 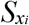 is the stoichiometric mass balance coefficients of the *i*th metabolite, and 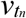 is a vector of the fluxes at timepoint *t_n_*. The kinetic equations (if known) of 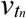 also contain relative abundances and response factors and are as shown in the inset (Michaelis-Menten kinetics are used as an example, where *V_maxf_* and 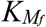 are kinetic parameters of the *f*th flux).

### Solving for flux distributions in the optimization approach

The only information that MetaboPAC assumes is known is the stoichiometry and metabolite-dependent allosteric regulation of the system, the kinetic structure of the system (if the kinetic equations approach is used), and the relative abundances of the data. In the optimization approach, the flux profiles of the reactions in the system are used to calculate some of the penalties that describe the relationship between inferred absolute concentrations and the reactions they control. Because fluxomics data are not assumed to be available, the fluxes must be inferred by solving Equation 1. As in the kinetic equations approach, the rate of change is determined by calculating the difference in relative abundance between two timepoints divided by the time difference. This rate of change is divided by the corresponding response factor to infer the rate of change of absolute concentration for each metabolite. While the fluxes of a determined system can be trivially calculated, underdetermined systems have an infinite number of flux solutions. To choose a single solution, the optimization approach uses the Moore-Penrose pseudoinverse, which minimizes the norm of the flux solution [26]. If the kinetic equations of some of the fluxes are known, they can be used to create a less underdetermined system that could possibly be determined or even overdetermined, which would allow a unique flux solution to be found.

### Noise-added data

To generate noisy data that more closely represent experimental metabolomics data, we used two sampling frequencies and two coefficients of variation (CoV) for randomly-added noise, for a total of four conditions. Sampling frequencies of 50 and 15 timepoints (nT) and CoVs of 0.05 and 0.15 were tested, where a higher CoV represents more noise (experimental error). Starting from the data generated by the ODEs defining the systems, each concentration value in each metabolomics dataset was replaced with a random value drawn from *N*_*i*,*k*_ ~ (*y*_*i*_(*t*_*k*_),*CoV*∙*y*_*i*_(*t*_*k*_)), where *y_i_*(*t_k_*) is the value of metabolite *i* at timepoint *k*. Noisy data was smoothed using a Gaussian filter with a window of one-fourth the length of the simulated time interval.

### Evaluation metrics and comparing to baseline methods

To measure the performance of MetaboPAC, we calculated the relative difference between the true and predicted values of the response factors using a logarithmic scale and determined if it was within a range of log_2_(1.1), log_2_(1.3), and log_2_(1.5) error, as shown in Equation 3. *RF_T_* is the true response factor, *RF_P_* is the predicted response factor, and *x* is the value that determine the log_2_ error range (i.e. 1.1, 1.3, or 1.5). We found that using absolute percent error instead of a logarithmic scale could lead to large error ranges that would make the evaluation metric less meaningful. For example, 100% error for a response factor of 500 would cover a range from 0 to 1000 (i.e. the entire search space of response factors for this work), whereas using a logarithmic scale would cover a range from only 250 to 1000. The percentages of predicted response factors that were within each log_2_ error range were compared among the methods assessed.

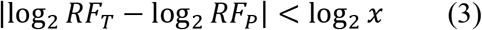

To provide a baseline performance for predicting response factors, we examined two other methods for predicting response factors that were compared to MetaboPAC. The first method randomly predicts response factors using a uniform distribution between 1 and 1000 for each metabolite. The second method uses a response factor of 500 for each metabolite, as predicted response factors close to the middle of the search space will have the greatest chance of being contained within the error range of the true response factor if the response factors are chosen from a uniform distribution.

## Results

### MetaboPAC performance on noiseless data

When initially assessing the performance of MetaboPAC on the two synthetic systems, we found the framework to perform exceptionally well on noiseless data (Figure 2). For all percentages of known kinetic equations, MetaboPAC performed significantly better than the random response factors and response factors of 500 for each of the log_2_ error ranges examined. For the underdetermined system with regulation, MetaboPAC performed significantly better than the other two methods when at least 60% of the kinetic equations were known for the log_2_(1.1) error range and when at least 40% of the kinetic equations were known for the log_2_(1.3) and log_2_(1.5) error ranges. Unsurprisingly, as the percentage of known kinetic equations increased, the accuracy of predicted response factors also generally increased for MetaboPAC, with 100% of the response factors within the log_2_(1.1) error range for both systems when 100% of the kinetic equations were known. As expected, the response factors predicted when using the kinetic equations approach were more accurate than the predictions by the optimization approach (Figure S2). Figure S3 shows the mean percentage of response factors predicted by either the kinetic equations approach or optimization approach across different percentages of known kinetic equations.

**Figure 2.**
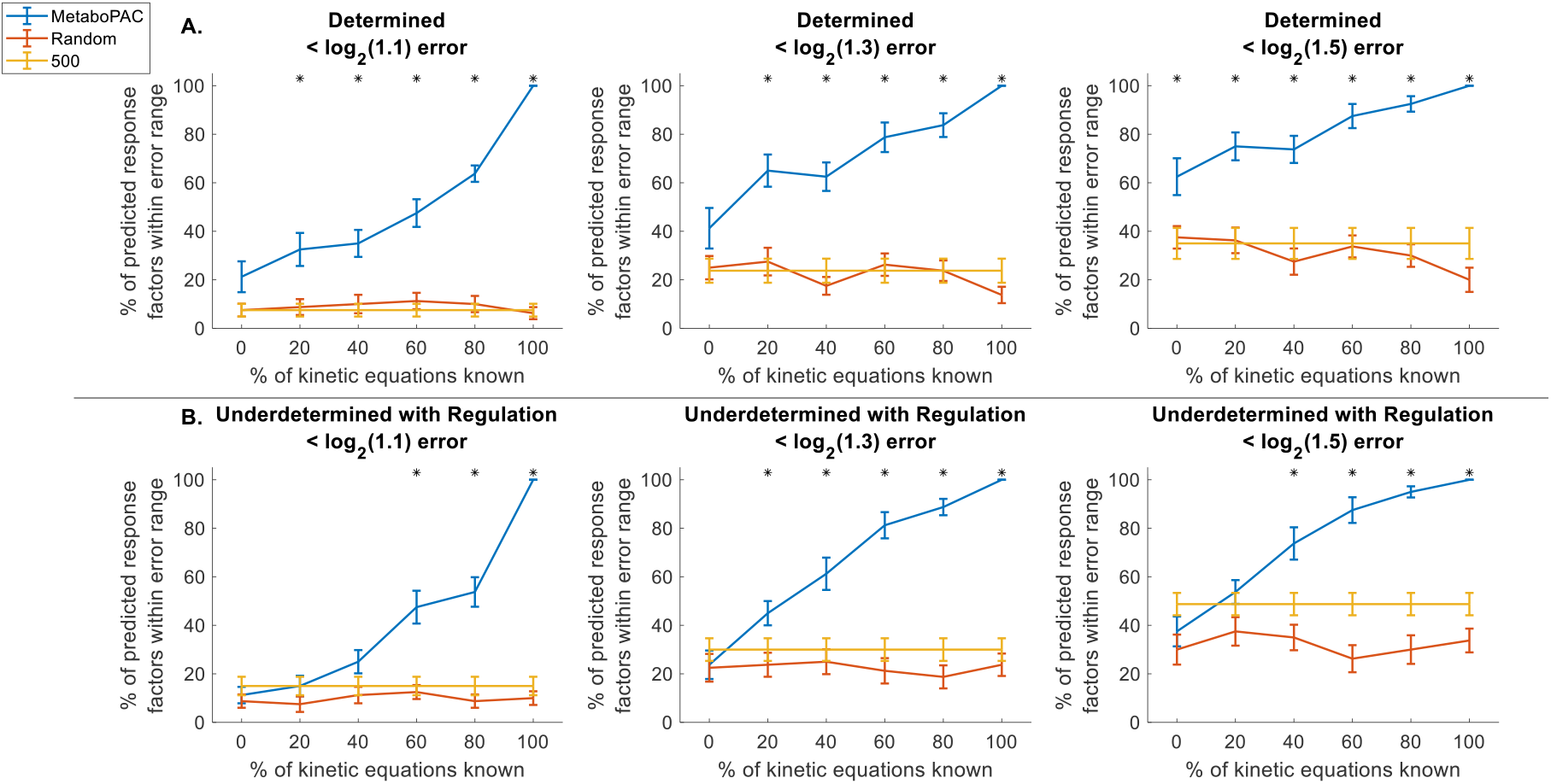
MetaboPAC performance on noiseless data for synthetic systems. MetaboPAC compared to random response factors and response factors of 500 for the A. determined and B. underdetermined with regulation systems using error ranges of log_2_(1.1), log_2_(1.3), and log_2_(1.5). Lines represent the mean percent of predicted response factors within the error ranges for each method. Error bars represent the standard error of the mean (n = 20 for different sets of true response factors). Asterisks denote when MetaboPAC performed significantly better at predicting response factors than both of the other two methods (two-sample t-test with α = 0.05).

Testing MetaboPAC on the two biological systems with noiseless data yielded similar results (Figure 3). For the *S. cerevisiae* system, MetaboPAC performed significantly better than the other two methods across all log_2_ error ranges when at least 40% of the kinetic equations were known, except for one case in the log_2_(1.5) error range. In the *E. coli* system, 60% of kinetic equations were required to be known for MetaboPAC to perform significantly better than the other two methods across all log_2_ error ranges. Once again, the kinetic equations approach typically outperformed the optimization approach (Figure S4). While the performance of MetaboPAC on the biological systems was not as high as the performance on the synthetic systems, it was still able to predict at least 58.9% of the response factors within log_2_(1.1) error and at least 83% of the response factors within log_2_(1.3) error in both systems when 100% of the kinetic equations are known.

**Figure 3.**
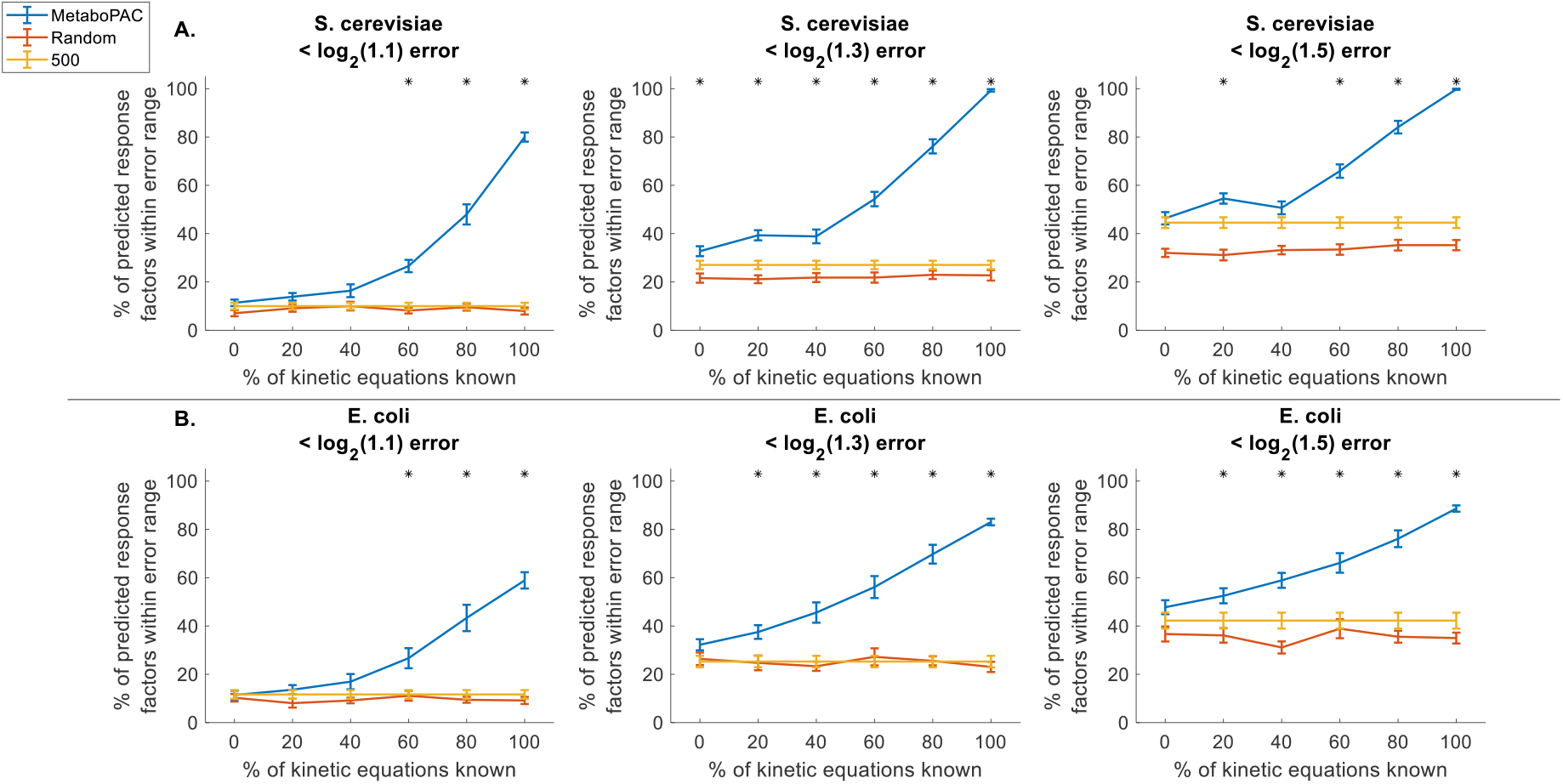
MetaboPAC performance on noiseless data for biological systems. MetaboPAC compared to random response factors and response factors of 500 for the A. *S. cerevisiae* and B. *E. coli* systems using error ranges of log_2_(1.1), log_2_(1.3), and log_2_(1.5). Lines represent the mean percent of predicted response factors within the error ranges for each method. Error bars represent the standard error of the mean (n = 20 for different sets of true response factors). Asterisks denote when MetaboPAC performed significantly better at predicting response factors than both of the other two methods (two-sample t-test with α = 0.05).

### MetaboPAC performance on noise-added data

While noiseless data provides a good benchmark for the performance of MetaboPAC under ideal conditions, real experimental metabolomics data will have some degree of noise. To test the robustness of MetaboPAC under more realistic conditions, we assessed MetaboPAC on datasets with different sampling frequencies (nT = 50 or 15) and different amounts of added noise (CoV = 0.05 or 0.15). In the synthetic systems, MetaboPAC was significantly better than both the random and 500 response factor approaches for almost all log_2_ error ranges (Figure 4 and Figure 5) when 100% of kinetic equations were known under the low sampling frequency and high noise condition (nT = 15, CoV = 015). We also determined that MetaboPAC generally performed the best in the conditions with low amounts of noise (CoV = 0.05) and often only required 60% or 80% of the kinetics to be known to outperform the other methods. As found in the noiseless condition, the accuracy of the kinetic equations approach was higher than the accuracy of the optimization approach in most cases (Figure S6 and Figure S7). While there was a predictable decrease in overall performance compared to the results when using noiseless data, MetaboPAC was still able to predict 56.3% and 100% of response factors within log_2_(1.5) error for the determined and underdetermined system with regulation, respectively, when 100% of the kinetic equations were known. Surprisingly, MetaboPAC seems to perform better on the underdetermined system with regulation compared to the determined system at 100% known kinetic equations, despite the increase in complexity. The simplicity of the mass balance equations in the determined system (with fewer reaction kinetics and no regulation, and therefore fewer instances of response factors within the mass balance equations) may actually hinder identification of accurate response factors in this instance.

**Figure 4.**
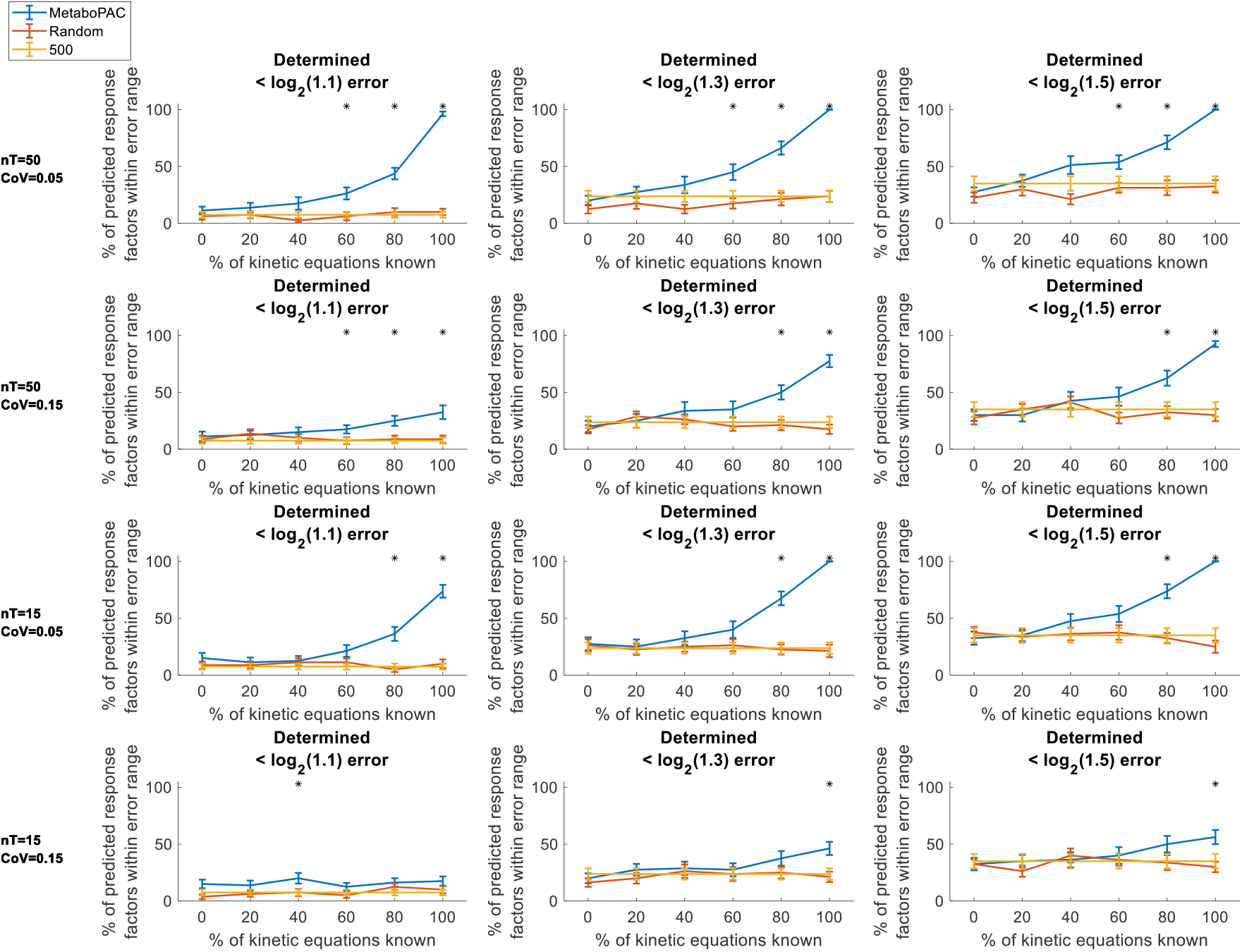
MetaboPAC performance on all conditions of noisy data for the determined system. MetaboPAC compared to random response factors and response factors of 500 for the determined system using error ranges of log2(1.1), log2(1.3), and log2(1.5) on data with different sampling frequencies (nT = 50 or 15) and noise added (CoV = 0.05 or 0.15). Lines represent the mean percent of predicted response factors within the error ranges for each method. Error bars represent the standard error of the mean (n = 20 for different sets of true response factors). Asterisks denote when MetaboPAC performed significantly better at predicting response factors than both of the other two methods (two-sample t-test with α = 0.05).

**Figure 5.**
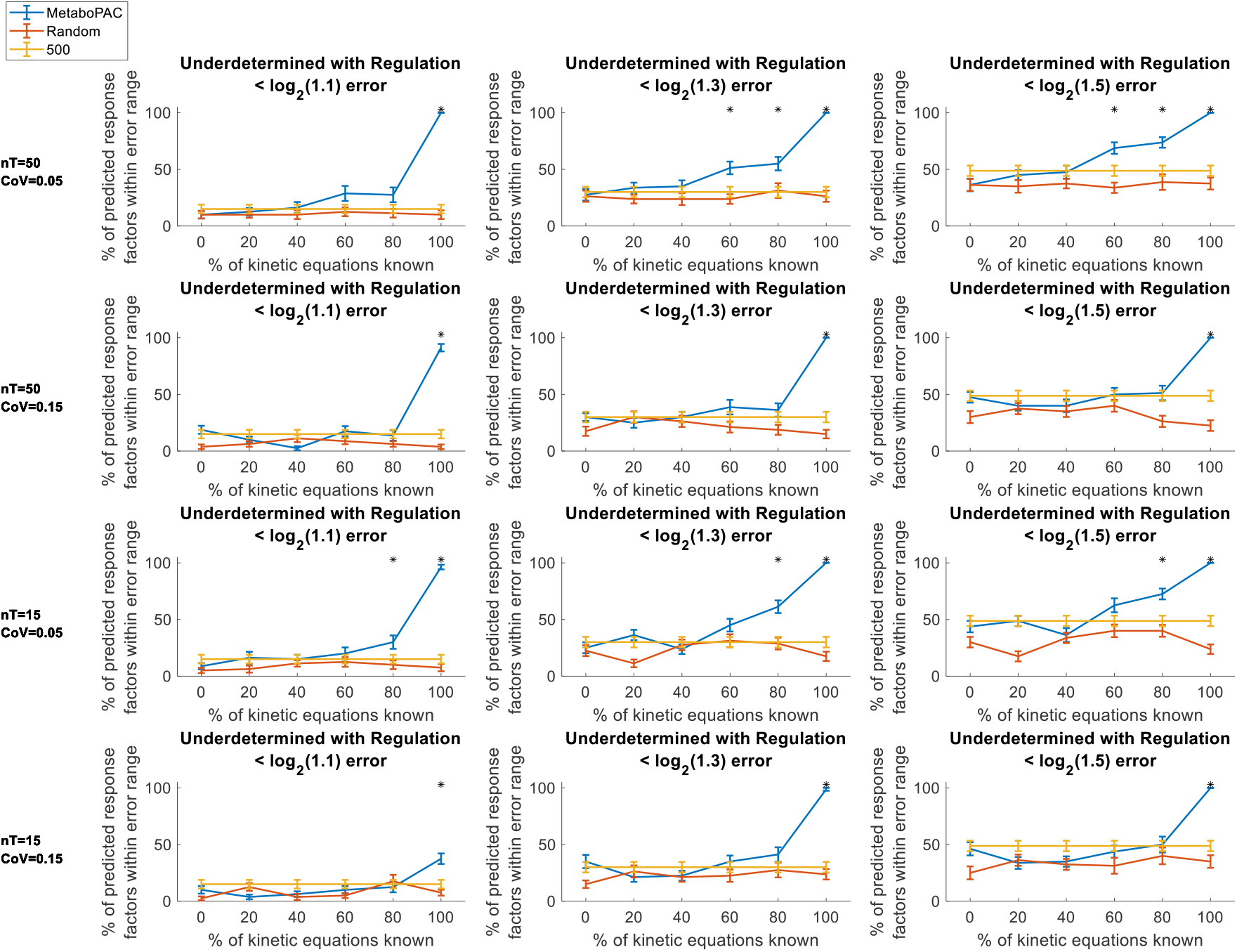
MetaboPAC performance on all conditions of noisy data for the undetermined system with regulation. MetaboPAC compared to random response factors and response factors of 500 for the underdetermined system with regulation using error ranges of log2(1.1), log2(1.3), and log2(1.5) on data with different sampling frequencies (nT = 50 or 15) and noise added (CoV = 0.05 or 0.15). Lines represent the mean percent of predicted response factors within the error ranges for each method. Error bars represent the standard error of the mean (n = 20 for different sets of true response factors). Asterisks denote when MetaboPAC performed significantly better at predicting response factors than both of the other two methods (two-sample t-test with α = 0.05).

For the two biological systems, MetaboPAC was still found to significantly predict response factors more accurately than random response factors or response factors of 500 for most of the log_2_ error ranges when 100% of the kinetic equations were known under low sampling and high noise conditions. Once again, MetaboPAC showed some improved performance when under conditions where there was a high sampling frequency or low noise (Figure 6 and Figure 7). Interestingly, the optimization approach often performed better than the kinetic equations approach when a low percentage of kinetic equations was known (Figure S8 and Figure S9), which was less common in the synthetic systems. In some cases, the optimization approach alone (0% known kinetic equations) was even significantly better than randomly predicting response factors or response factors of 500. This observation illustrates that both the kinetic equations and optimization approaches are important to the framework. Overall, MetaboPAC demonstrates that for biological system where the kinetic rate laws of metabolic pathways are known *a priori*, it can offer useful information for inferring some absolute concentrations.

**Figure 6.**
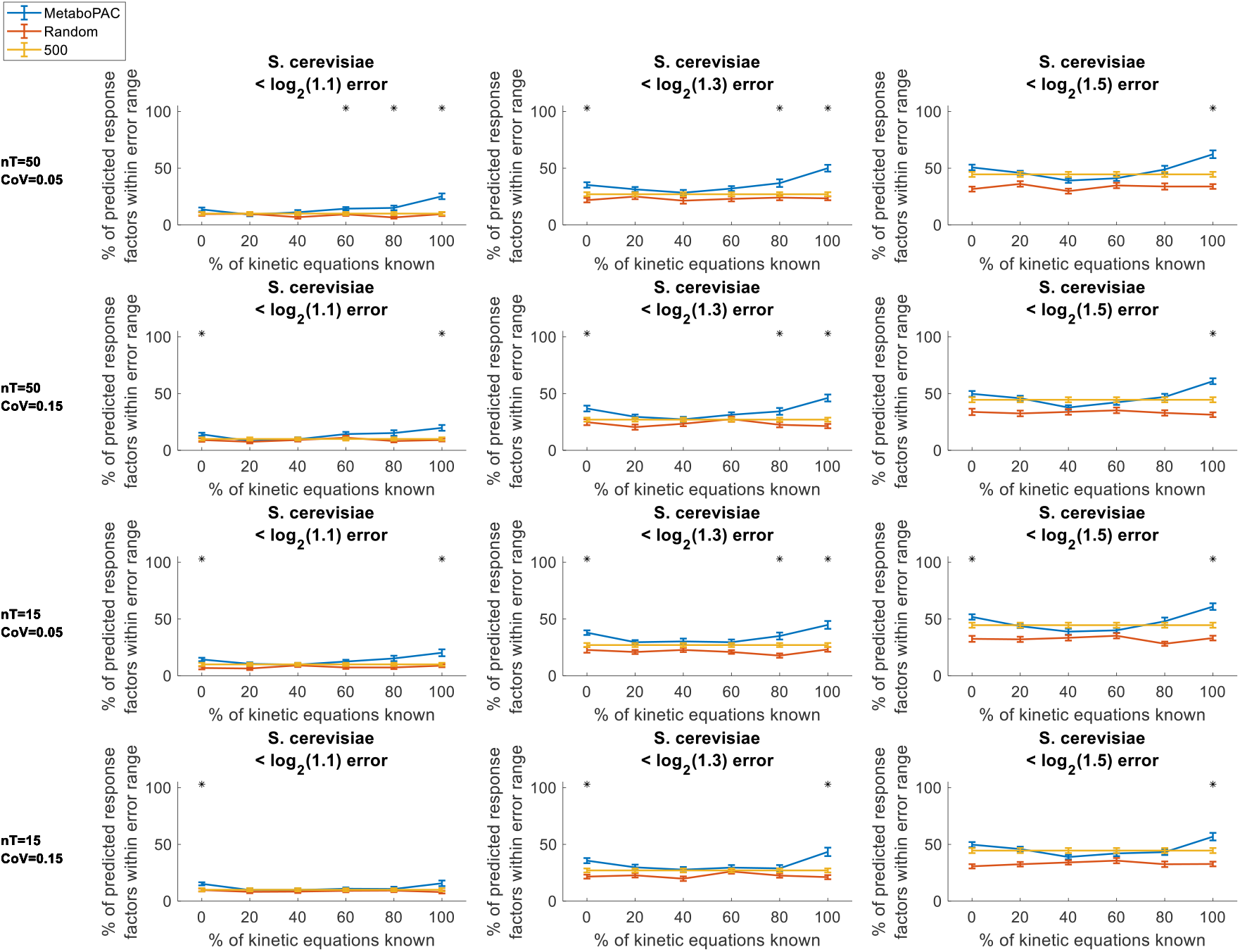
MetaboPAC performance on all conditions of noisy data for the *S. cerevisiae* system. MetaboPAC compared to random response factors and response factors of 500 for the S. cerevisiae system using error ranges of log2(1.1), log2(1.3), and log2(1.5) on data with different sampling frequencies (nT = 50 or 15) and noise added (CoV = 0.05 or 0.15). Lines represent the mean percent of predicted response factors within the error ranges for each method. Error bars represent the standard error of the mean (n = 20 for different sets of true response factors). Asterisks denote when MetaboPAC performed significantly better at predicting response factors than both of the other two methods (two-sample t-test with α = 0.05).

**Figure 7.**
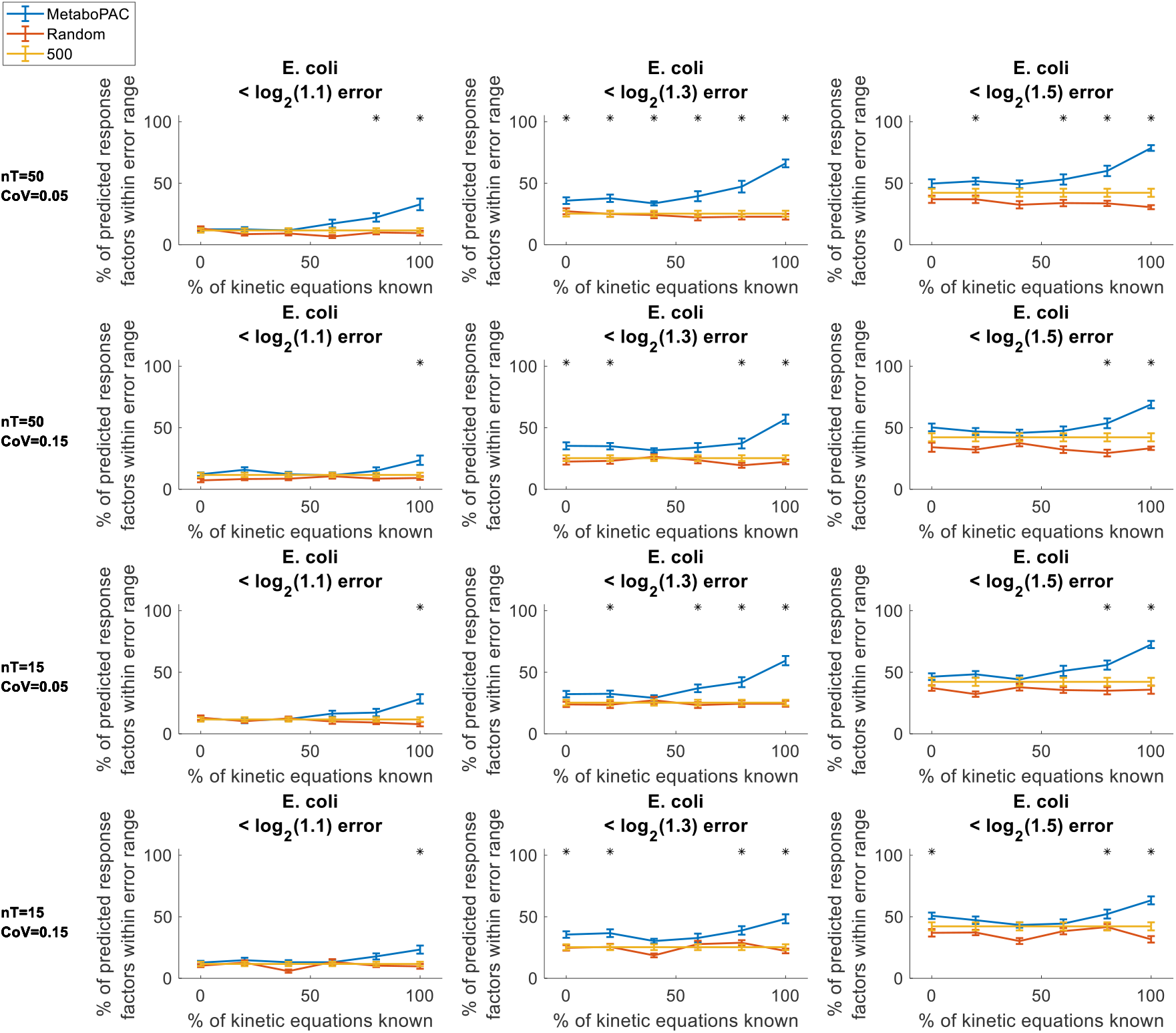
MetaboPAC performance on all conditions of noisy data for the *E. coli* system. MetaboPAC compared to random response factors and response factors of 500 for the E. coli system using error ranges of log2(1.1), log2(1.3), and log2(1.5) on data with different sampling frequencies (nT = 50 or 15) and noise added (CoV = 0.05 or 0.15). Lines represent the mean percent of predicted response factors within the error ranges for each method. Error bars represent the standard error of the mean (n = 20 for different sets of true response factors). Asterisks denote when MetaboPAC performed significantly better at predicting response factors than both of the other two methods (two-sample t-test with α = 0.05).

## Discussion

MetaboPAC has shown substantial potential to provide accurate absolute concentrations for metabolites in well-studied metabolic pathways (e.g. central carbon metabolism) whose kinetics have been previously determined for various biological systems. Metabolomics research in common microorganisms, such as *E. coli* and *S. cerevisiae*, could benefit significantly from MetaboPAC, as it will allow metabolomics data to be more seamlessly integrated with metabolic modelling frameworks and data analysis methods that require absolute concentrations. The key component of MetaboPAC is the use of mass balances within a system with known stoichiometry. Previously, mass balances have been used to determine quenching leakage in metabolomics [27], but to the best of our knowledge, this is the first time mass balances have been used in the context of inferring absolute concentrations. Because MetaboPAC leverages the mass balances of a system to predict its response factors, it is unsurprising that the performance of MetaboPAC is hindered under conditions with high noise, as the mass balances can be affected. Nevertheless, MetaboPAC still significantly outperformed the other two methods assessed when all kinetic equations are known, suggesting that systems with known kinetic structures would benefit from MetaboPAC.

One of the strengths of MetaboPAC is that additional information can be easily integrated into the framework to reduce the number of possible sets of response factors. If the minimum or maximum possible or predicted concentrations of each (or a few) metabolites are known, this can greatly reduce the search space of possible sets of response factors. We found that constraining the range of possible response factors of one metabolite would often lead to the range of possible response factors of other metabolites also being constrained, especially metabolites nearby in the metabolic pathway. This is likely due to their mass balances sharing some of the same reaction fluxes. Along those lines, chemical standards could be used for some metabolites, which would decrease the number of response factors MetaboPAC would need to predict and would once again lead to more constrained ranges of possible response factors for some metabolites.

Along with incorporating additional information directly into the framework, MetaboPAC could also be used concurrently with other metabolomics methods to improve the accuracy of response factor predictions. The use of methods focused on predicting ionization efficiencies [20–22] in conjunction with MetaboPAC could provide further insight about which response factors predicted by MetaboPAC are most likely to be accurate if the inferred concentrations of MetaboPAC and the ionization efficiency-based platforms are similar. When working with underdetermined systems, methods such as dynamic flux estimation [28, 29] or flux balance analysis [30] could be applied to determine more likely flux distributions than the Moore-Penrose pseudoinverse approach used in MetaboPAC, which could lead to improved predictions of response factors when using the optimization approach. Alternatively, if minimum or maximum values of individual fluxes are known, this could also reduce the number of possible flux solutions and benefit the calculation of penalties in the optimization approach.

In this proof-of-principle work, there are two key assumptions we have used to initially assess MetaboPAC. These assumptions are reasonable to assume for this work, but they will need to be adjusted in the future for MetaboPAC to be more widely applicable. First, we assumed the relationship between relative abundances and their absolute concentrations is linear. These relationships are not always linear in experimental data and non-linear relationships will need to be considered in the future. Both the kinetic equations and optimization approaches within MetaboPAC can be easily adjusted to account for non-linear relationships, but determining which metabolites have non-linear relationships is a more difficult problem and will need to be explored if this information is not known *a priori*.

MetaboPAC also assumes the true response factors are sampled from a uniform distribution. Under the most realistic conditions, this may not be the case. To further assess the robustness of MetaboPAC, we tested the framework on noiseless relative abundance data from the two biological systems (assuming all kinetic equations are known) where the true response factors were drawn from a log uniform distribution (Figure S10). While there is a decrease in performance when using MetaboPAC on both biological systems compared to the results in Figure 3, this drop in performance is also seen in both the random and 500 response factor methods. MetaboPAC is still significantly better than the other two methods by a wide margin in all log_2_ error ranges examined, which indicates that MetaboPAC is still suitable even if response factors are not uniformly distributed.

While the results presented here are promising, there are MetaboPAC currently has some limitations. First, because MetaboPAC leverages the stoichiometric mass balance of a biological system to identify response factors, it can only be used in the context of cellular metabolism in its current form. For example, MetaboPAC could not infer the absolute concentrations of metabolites in a blood sample because blood metabolite profiles are determined from metabolic contributions from across organs or systems in an organism with no stoichiometric basis to connect concentrations [31].

Perhaps the greatest obstacle for MetaboPAC is noisy data. In our results, we have determined that MetaboPAC performs particularly well on datasets with little or no noise, even when the kinetics of the system are not fully known. Under the noisiest condition (CoV = 0.15), we still generally find MetaboPAC to perform significantly better than other methods for predicting response factors when all kinetic equations were known, but there is a noticeable decrease in accuracy. Leveraging the mass balances of a metabolic system is one of the critical ideas behind MetaboPAC and it is not surprising that data with high noise affect these calculations. Here, we have used a Gaussian filter to smooth the noisy data, which has proven to be effective, but other venues for mitigating the effect of noise should be considered. Exploring other options to reduce noise, such as filtering, normalization, scaling, other smoothing methods [32–34], or using triplicate samples as is common when collecting metabolomics data, could prove useful and lead to an increase in performance.

## Conclusion

The need for chemical standards in mass spectrometry methods to absolutely quantify metabolomics data has been a challenging obstacle that has prevented the direct use of metabolomics in many metabolic modelling tools. MetaboPAC is a critical step toward preprocessing metabolomics data so that it can be readily used with metabolic modelling and other computational platforms that require absolute concentrations of metabolites. For well-studied systems, where the entire kinetic structure is known, MetaboPAC can infer absolute concentrations with high accuracy. Under conditions where the amount of noise in the data is minimal, MetaboPAC can still provide valuable information if the kinetic equations are only partially known. As research in metabolomics continues to grow and more computational frameworks aim to harness all the information that metabolomics data has to offer, MetaboPAC has the potential to become a powerful tool for absolute quantification in metabolomics.

