## Supplementary for "Leveraging the mass balances of cellular metabolism to infer absolute concentrations from relative abundance metabolomics data"

### Supplementary Information

**Table S1: Penalties used in the optimization approach of MetaboPAC**

| Penalty | Description | Reasoning |
|---|---|---|
| Mass balance | Calculate the sum of squared residuals between the inferred change in absolute concentration over time calculated from the raw relative abundance data (i.e. the change in relative abundance over time divided by the predicted response factor) and the inferred change in absolute concentration over time calculated from the stoichiometry of the system and inferred fluxes (i.e. Equation 1 in the Methods). | If the change in absolute concentration over time is vastly different between the two calculations (i.e. the sum of squared residuals is greater than zero), the predicted response factors have failed to produce inferred absolute concentration and flux profiles that do not violate any mass balances in the system. |
| Maximum concentration | If the inferred absolute concentration for any metabolite is above 5 mM or 50 mM for synthetic and biological systems, respectively, add a penalty equal to the maximum value of all inferred concentrations. | It is reasonable to assume that for many metabolites, there can be a general estimate for a maximum concentration that is biologically feasible, either due to limits in production or cell toxicity. Here, we use a single threshold for all metabolites, but imposing individual maximum thresholds would lead to better response factor predictions. |

| | | |
|---|---|---|
| Correlation for mass action reaction with a single substrate | Calculate the correlation between the controller metabolite and inferred target flux. The correlation is expected to be positive (because metabolites induce mass action reactions), the penalty for each one-controller metabolite reaction equals the calculated correlation minus one. | If a reaction is only controlled by a single metabolite, the reaction rate should either increase or decrease as the concentration of the metabolite increases (assuming the kinetics of the reaction do not exhibit any behavior similar to substrate inhibition). |
| Curve fit for mass action reaction with a single substrate | Calculate the fit of a second-order polynomial to the controller metabolite and target flux data. The penalty for each one-controller metabolite reaction equals one minus the adjusted $R^2$ of the fit (adjusted for the number of coefficients). | A second-order polynomial should fit the data reasonably well if a reaction is controlled by a single metabolite (e.g. if the data is well-modeled by a Michaelis-Menten saturation curve). |
| Correlation for reactions regulated by two controller metabolites | Plot the data of one of the controller metabolites (x-axis) against the data of the inferred target flux (y-axis). Next, plot 23 vertical lines that are evenly spaced within the range of the controller metabolite that represent 23 constant concentrations. For one vertical line, identify if and where the line intersects with the data using the InterX function and linearly interpolate flux data at these intersection points using the closest higher and lower concentrations with sampled flux data. Calculate the Spearman correlation between the second controller metabolite and interpolated target flux at these intersection points where the first controller metabolite is constant. Repeat for each of the 23 vertical lines and then | The correlation between one controller metabolite and its target flux should be consistently close to +1 (activation) or -1 (inhibition) for any constant concentration value for the second metabolite. This assumes that a controller metabolite cannot switch from inducing to inhibiting a reaction (or vice versa) at high concentrations, such as in substrate inhibition. |

| | | |
|---|---|---|
| | calculate the mean correlation. To calculate the penalty, take the mean correlation and subtract (for metabolites expected to induce the reaction) or add (for metabolites expected to inhibit the reaction) one. Switch which metabolite is held constant and repeat the process. Finally, sum all penalties together. | |
| Fit to BST kinetic equations | For each reaction in a system, fit the inferred absolute concentration and flux data to a BST equation [1] representing the reaction rate. Calculate the sum of squared residuals of the fit. | A generic BST kinetic equation should fit reasonably well to correctly inferred absolute concentration and flux data. |

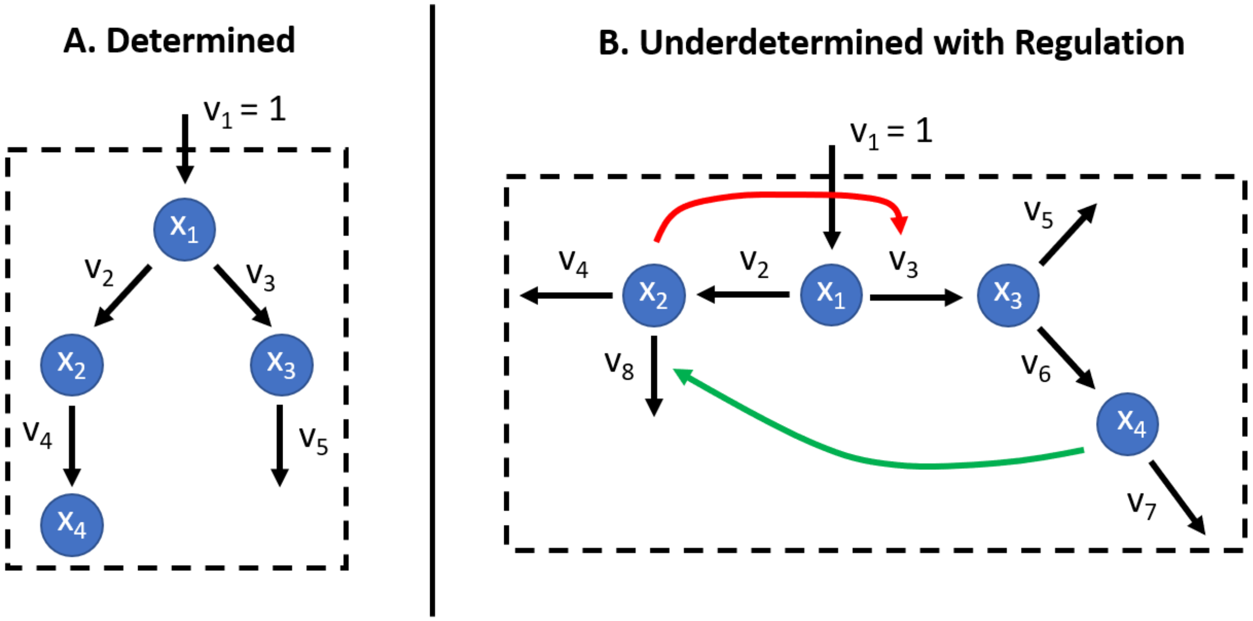


**Figure S1. Synthetic systems tested with MetaboPAC.** We built one determined synthetic system and one underdetermined synthetic system with regulation using Michaelis-Menten kinetics for each reaction. $x_i$ represents the $i$th metabolite and $v_j$ represents the $j$th flux. In both systems, flux $v_1$ is assumed to be constant and known.

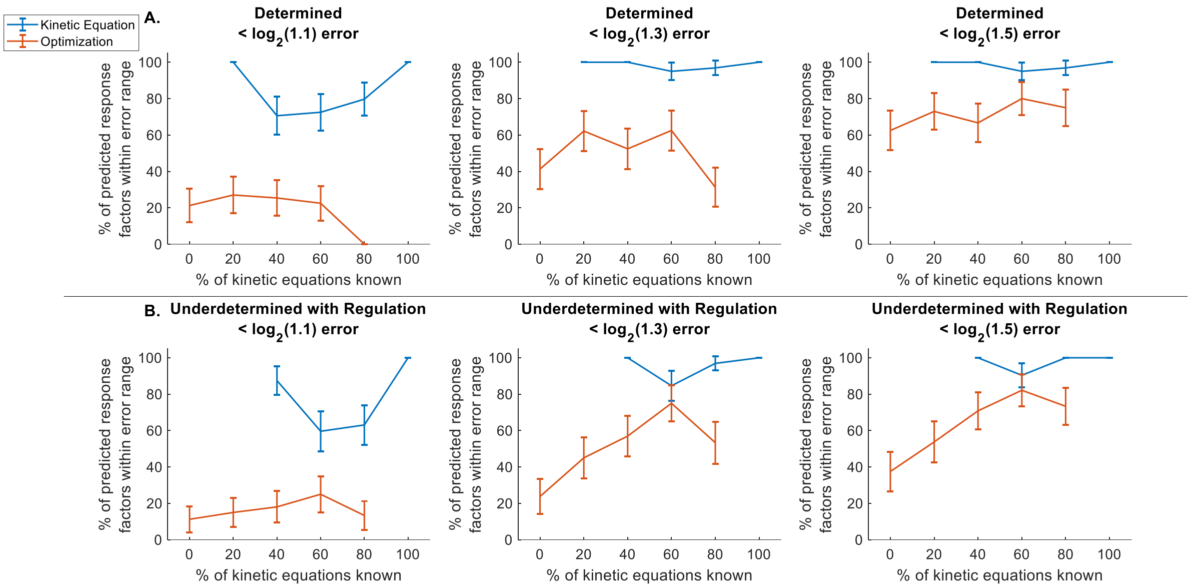


**Figure S2. Percent of response factors predicted by the kinetic equation and optimization approaches within each $\log_2$ error range for the synthetic systems when using noiseless data.** The kinetic equation approach generally predicted more accurate response factors than the optimization approach. Error bars represent the standard error of the mean (number of samples varies based on the percentage of kinetic equations known (Figure S3))**.**

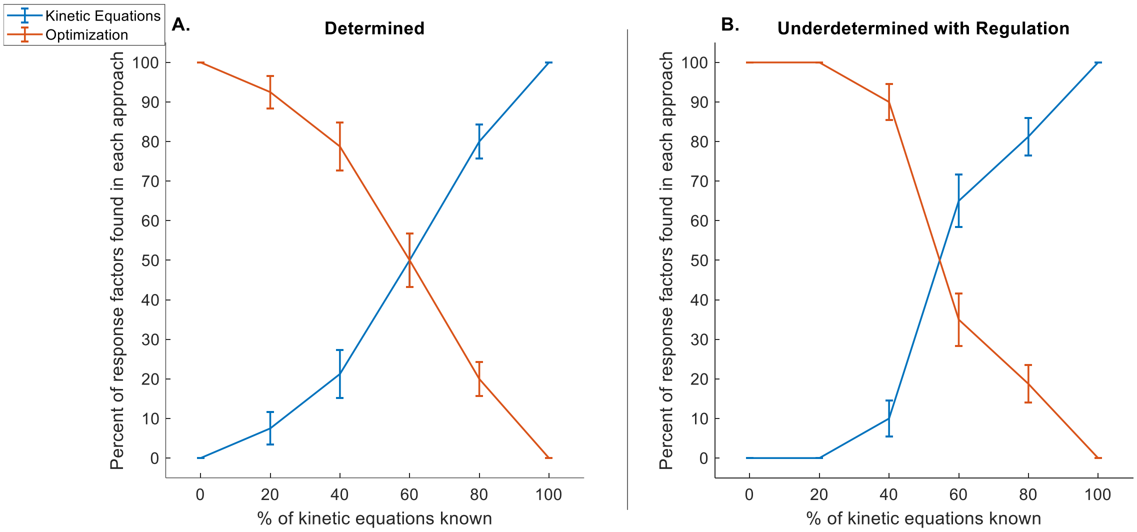


**Figure S3. Percentage of response factors predicted by the kinetic equation and optimization approaches in the synthetic systems.** As the percentage of known kinetic equations increases, it is more likely for response factors to be solvable using the kinetic equation approach. Error bars represent the standard error of the mean (number of samples varies based on the percentage of kinetic equations known).

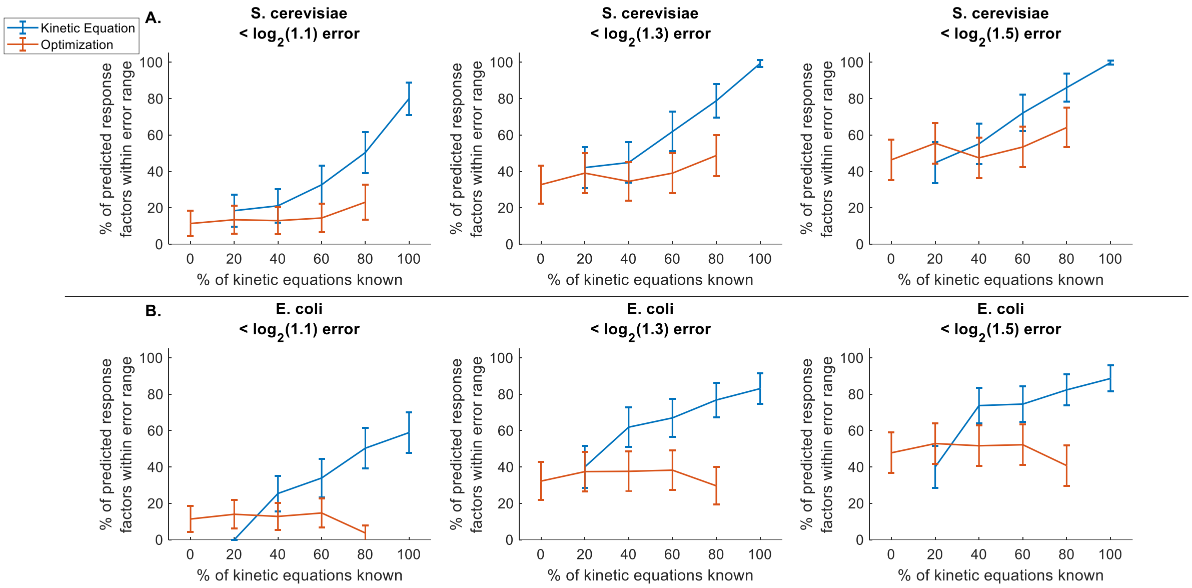


**Figure S4. Percent of response factors predicted by the kinetic equation and optimization approaches within each $log_2$ error range for the *S. cerevisiae* and *E. coli* systems when using noiseless data.** The kinetic equation approach generally predicted more accurate response factors than the optimization approach. Error bars represent the standard error of the mean (number of samples varies based on the percentage of kinetic equations known (Figure S5))**.**

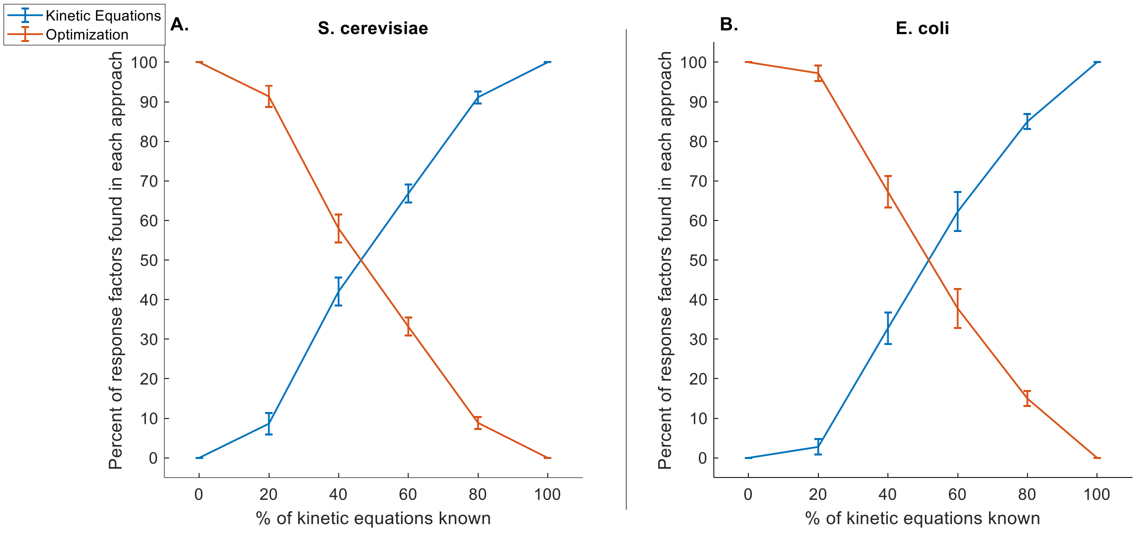


**Figure S5. Percentage of response factors predicted by the kinetic equation and optimization approaches in the *S. cerevisiae* and *E. coli* systems.** As the percentage of known kinetic equations increases, it is more likely for response factors to be solved using the kinetic equation approach. Error bars represent the standard error of the mean (number of samples varies based on the percentage of kinetic equations known).

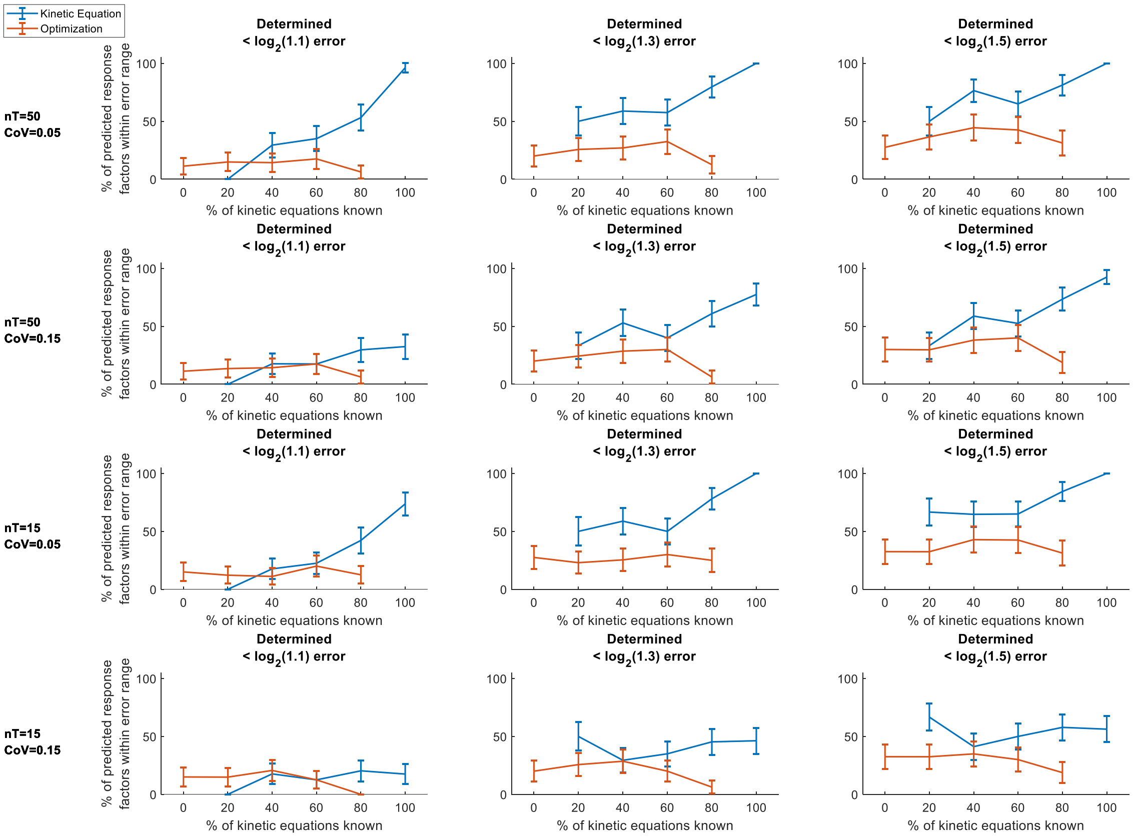


**Figure S6. Percent of response factors predicted by the kinetic equation and optimization approaches within each $\log_2$ error range for the determined system when using noisy data.** The kinetic equation approach generally predicted more accurate response factors than the optimization approach. Error bars represent the standard error of the mean (number of samples varies based on the percentage of kinetic equations known (Figure S3)).

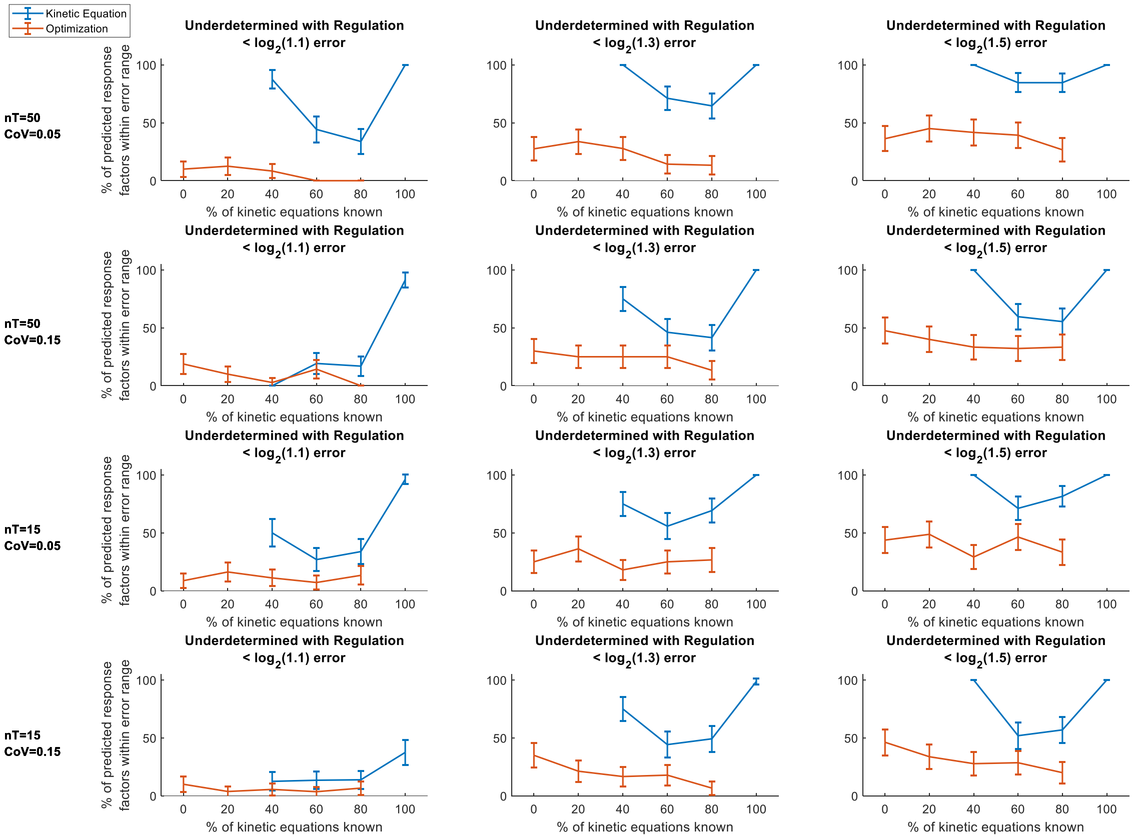


**Figure S7. Percent of response factors predicted by the kinetic equation and optimization approaches within each log₂ error range for the underdetermined system with regulation when using noisy data.** The kinetic equation approach generally predicted more accurate response factors than the optimization approach. Error bars represent the standard error of the mean (number of samples varies based on the percentage of kinetic equations known (Figure S3)).

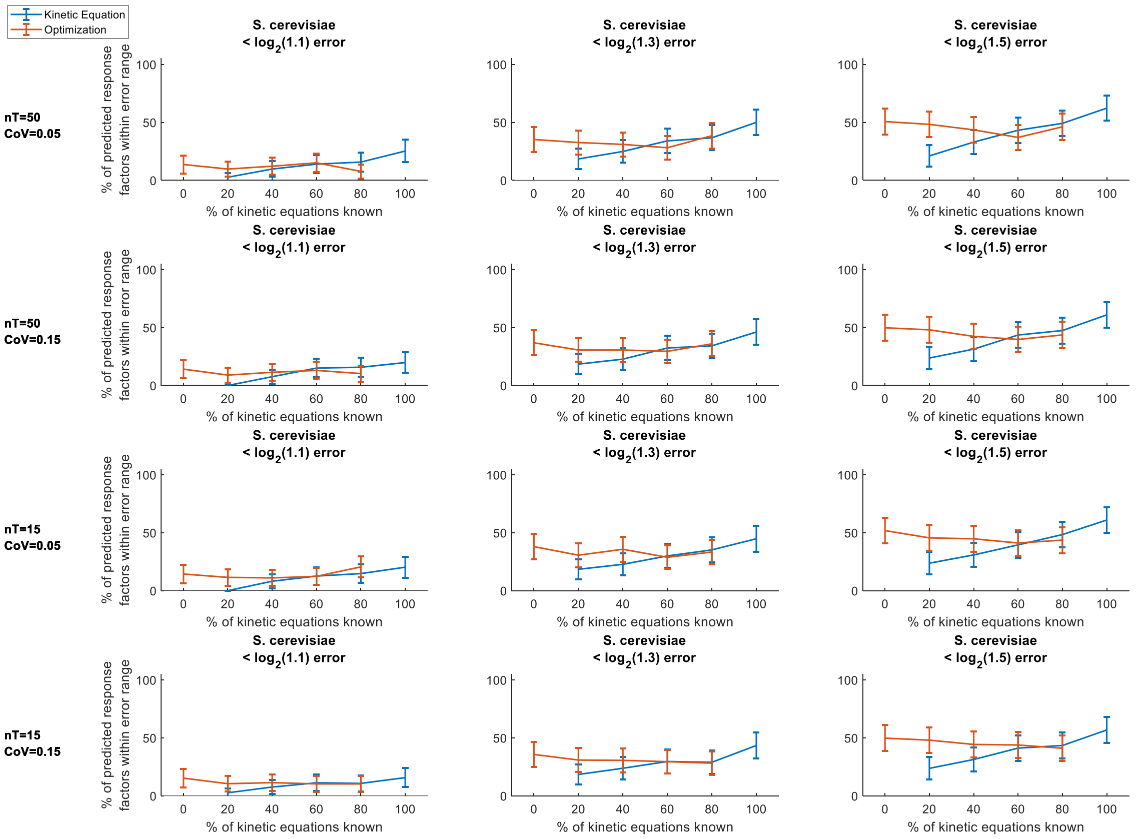


**Figure S8. Percent of response factors predicted by the kinetic equation and optimization approaches within each $\log_2$ error range for the *S. cerevisiae* system when using noisy data.** At low percentages of known kinetic equations, the optimization approach often performed better than the kinetic equation approach. At around 80% known kinetic equations, the kinetic equation approach began to have improved performance over the optimization approach. Error bars represent the standard error of the mean (number of samples varies based on the percentage of kinetic equations known (Figure S5)).

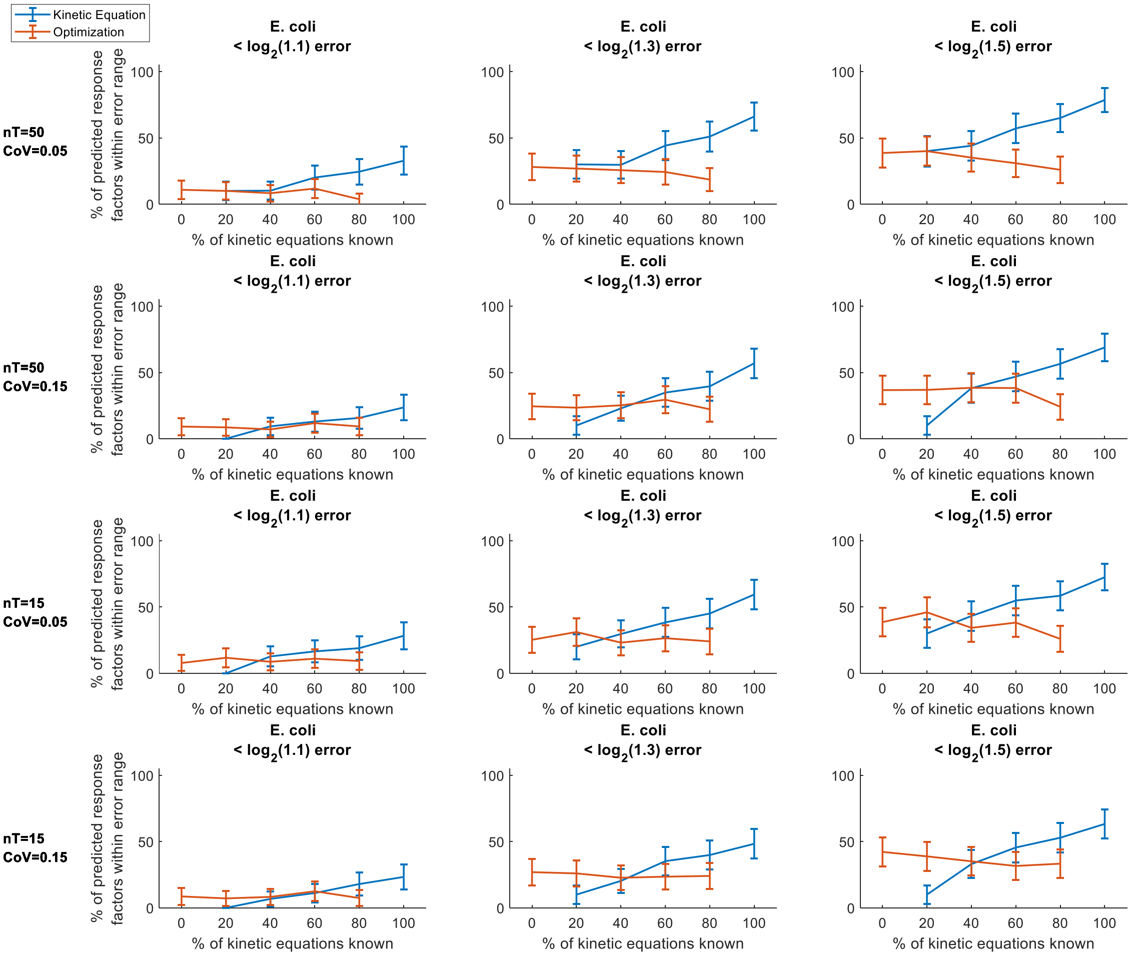


**Figure S9. Percent of response factors predicted by the kinetic equation and optimization approaches within each log$_2$ error range for the *E. coli* system when using noisy data.** At low percentages of known kinetic equations, the optimization approach often performed better than the kinetic equation approach. At around 60% known kinetic equations, the kinetic equation approach began to have improved performance over the optimization approach. Error bars represent the standard error of the mean (number of samples varies based on the percentage of kinetic equations known (Figure S5)).

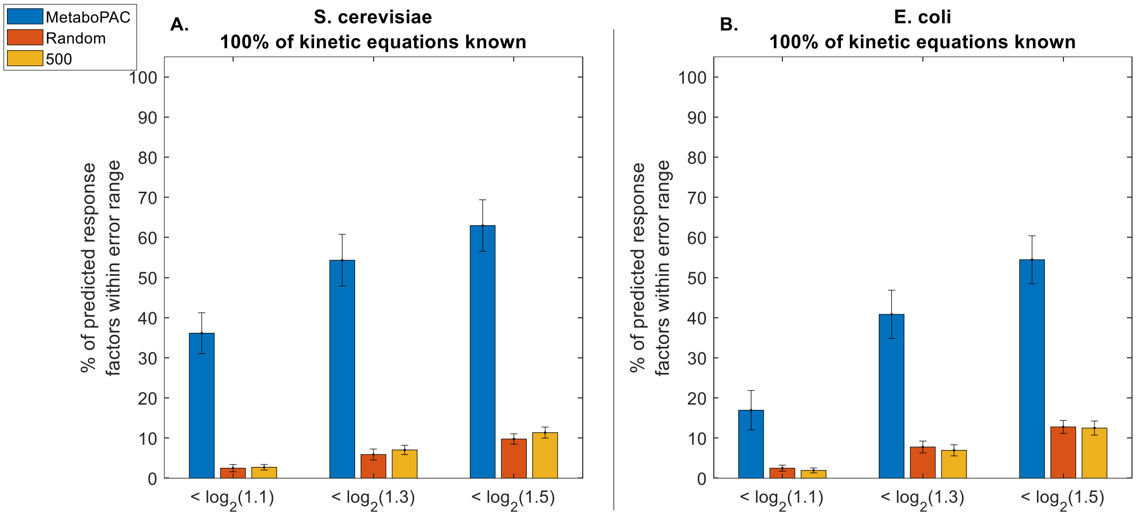


**Figure S10. MetaboPAC Performance when true response factors are sampled from a log uniform distribution.** To further test the robustness of MetaboPAC, the true response factors were drawn from a log uniform distribution instead of a uniform distribution for the two biological systems with 100% known kinetic equations and noiseless data. MetaboPAC still outperformed the other two methods across all $\log_2$ error ranges in both the *S. cerevisiae* and *E. coli* systems. Bars represent the mean percent of predicted response factors within the error ranges for each method. Error bars represent the standard error of the mean (n = 20 for different sets of true response factors).
